## Supplementary Information for "Breaking Barriers in Crosslinking Mass Spectrometry: Enhanced Throughput and Sensitivity with the Orbitrap Astral Mass Analyzer"

### Tabel of Figures

### Tabel of Tables

**Table S1**: Column parameters for PepMap and Aurora analytical column used in this study. .............................. 6
**Table S2**: Search parameters for linear and crosslink search. Parameters not listed here were left at default settings. ................................................................................................................................................ 8

### Supplemental Figures

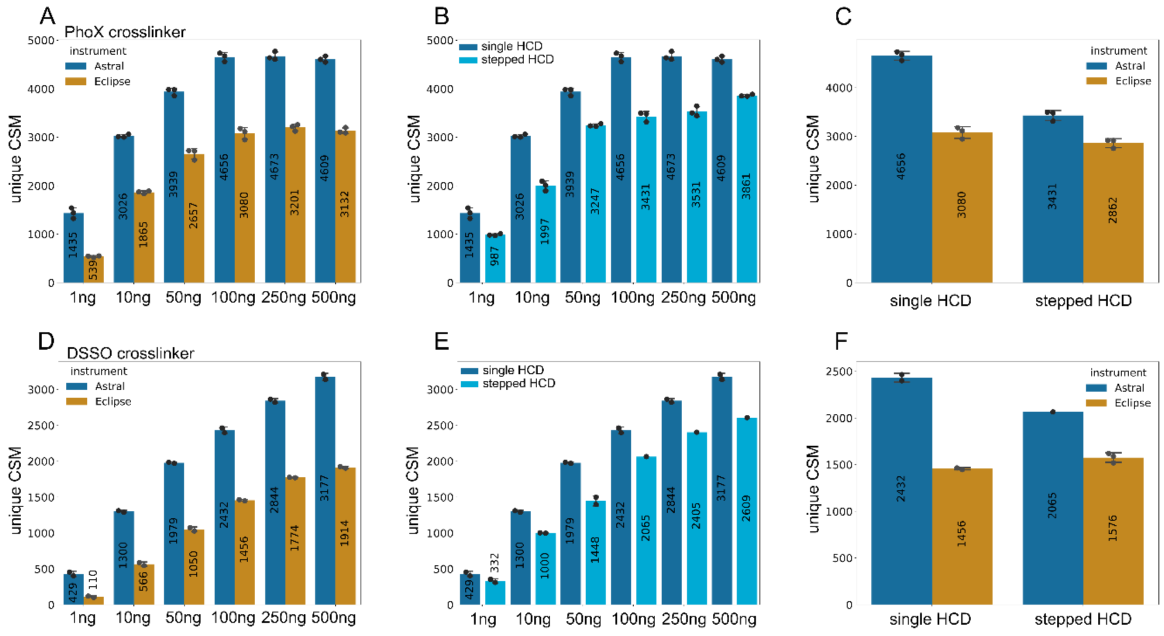

*Figure S1: Comparison of Astral and Eclipse unique crosslink spectrum match (CSM, 1% CSM level FDR) data with single and stepped HCD for PhoX (non-cleavable) and DSSO (cleavable) crosslinker. A: Dilution series of Cas9 crosslinked with PhoX starting from 1ng to 500 ng acquired on Eclipse and Astral in triplicates. The maximum number of CMSs is reached at 100 ng and plateauing afterwards. The Astral mass analyser (blue) outperforms the Eclipse (orange) data by more than 30%. B: Comparison between single HCD (dark blue) and stepped HCD (light blue) of the PhoX dilution series. Single HCD shows higher numbers of CSMs as stepped HCD for all injections amounts. C: Direct view on the difference between stepped and single HCD acquired on both instruments for 100 ng. The difference between single HCD and stepped HCD is dominant for Astral data (blue) but not significantly different for Eclipse data (orange). D: Dilution series of Cas9 crosslinked with DSSO starting from 1ng to 500 ng acquired on Eclipse and Astral in duplicates. The maximum number of unique crosslinks is reached with 500 ng. The Astral mass analyser (blue) outperforms the Eclipse (orange) data by more than 30% also for cleavable crosslinkers like DSSO. E: Comparison between single HCD (dark blue) and stepped HCD (light blue) of the DSSO dilution series. Single HCD shows also increased numbers of CSMs as stepped HCD for the DSSO dilution series. F: Direct view on the difference between stepped and single HCD acquired on both instruments for 100 ng. The difference between single HCD and stepped HCD is dominant for Astral data (blue) but not significantly different for Eclipse data (orange).*

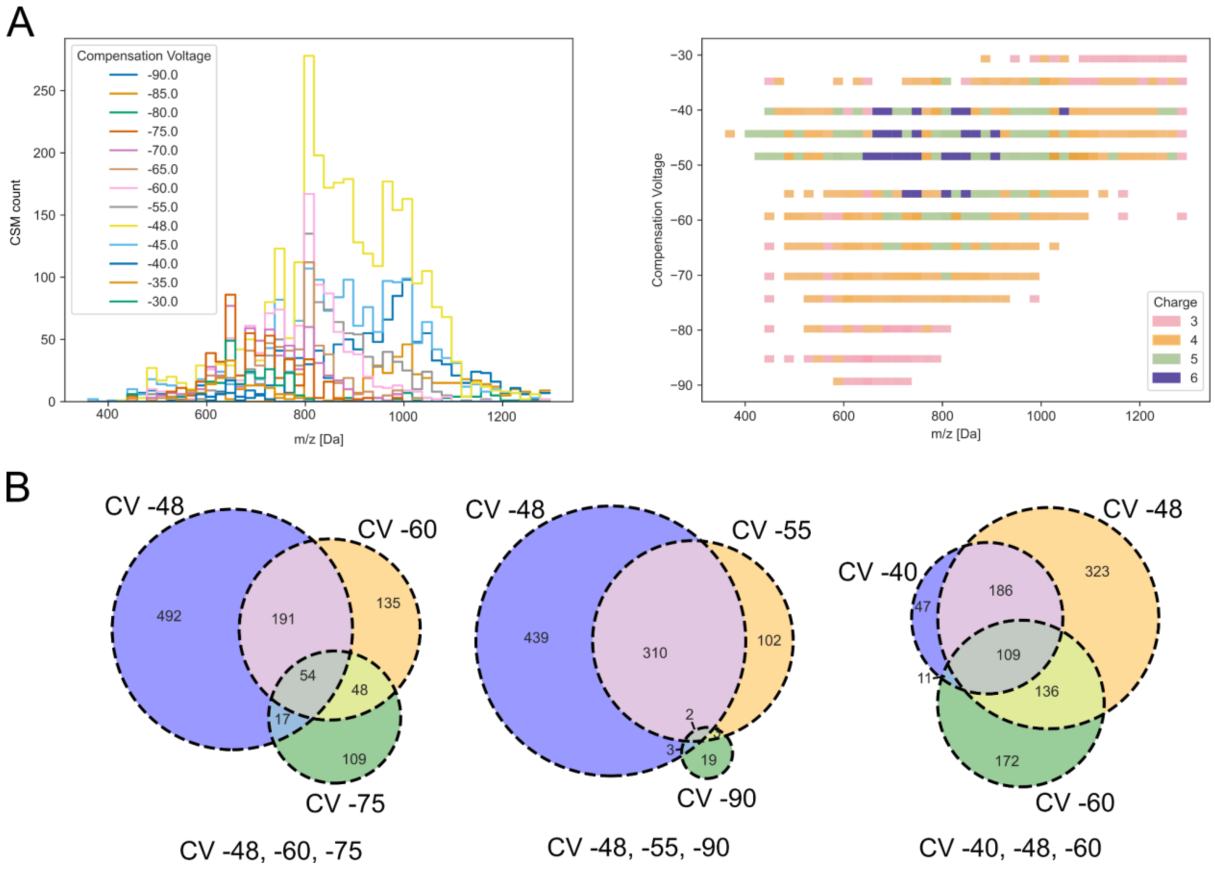

*Figure S2: Detailed view on differences between single CV measurements on CSM level to determine optimal CV combinations. A left: Histogram of each CV measurement concerning their m/z values. Single CVs are coloured according to the legend in 13 different colours. -48 V (yellow) shows a broad distribution across the whole m/z range with the maximum count of CSMs, while -90 V (dark blue) shows the minimum coverage with the least CSMs count. A right: CVs across the whole mass range coloured by charge states of identified CSMs. Charge 6 could be only identified from CV -40 V to -55 V. Charge 3 and 4 show a distribution across all CV values, while Charge 5 is only present in CV -35 V to -70 V. B: Based on an upset plot of all CV measurements, the three most promising CV combinations were selected for further evaluation. CV -48 V, -60 V, -75 V were selected to achieve the highest number of crosslinks with the least overlap, CV -48 V, -55 V, -90 V were selected to have the least overall overlap and CV 40 V, -48 V, -60 V to have the highest overlap and were used as common CV values for in-house crosslinking measurements. The unique crosslinks plotted in Venn diagrams have been summed up from CSMs with 1% CSM level FDR therefore the total number of links differs from Figure S3 where the FDR was calculated directly with 1% link level FDR.*

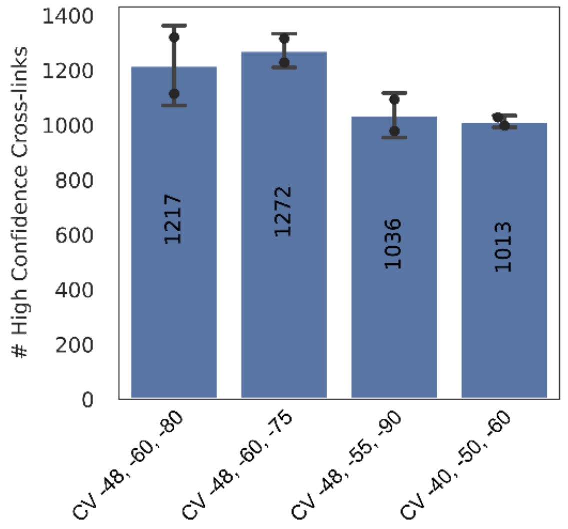

*Figure S3: Results of the CV optimization and combinatory experiment on unique crosslink level with 1% link level FDR. CV -48 V, -60 V, - 75 V gave the best results as expected. CV -48 V, -55 V, -90 V resulted in 19% fewer identifications, while CV -40 V, -50 V, -60 V gave 20% fewer unique crosslinks. CV -48 V, -60 V, -80 V are used as the QC acquisition method, with no significant change to the best-performing method (4% less identifications).*

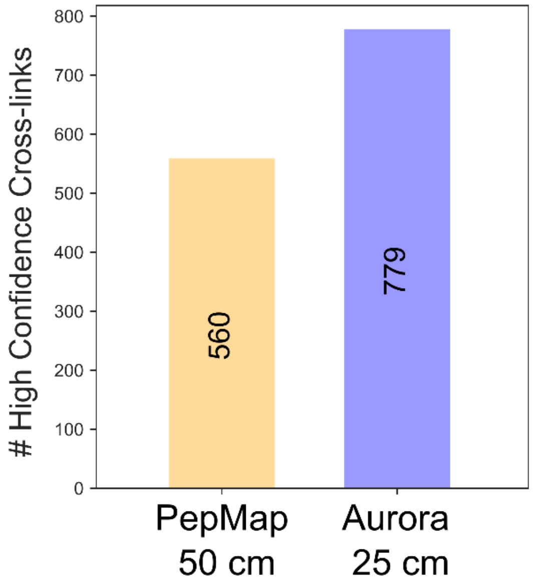

*Figure S4: Column comparison of PepMap 50 cm vs. Aurora Ultimate 25 cm analytical columns. Unique crosslink numbers of both columns with Aurora outperforming PepMap by 28%.*

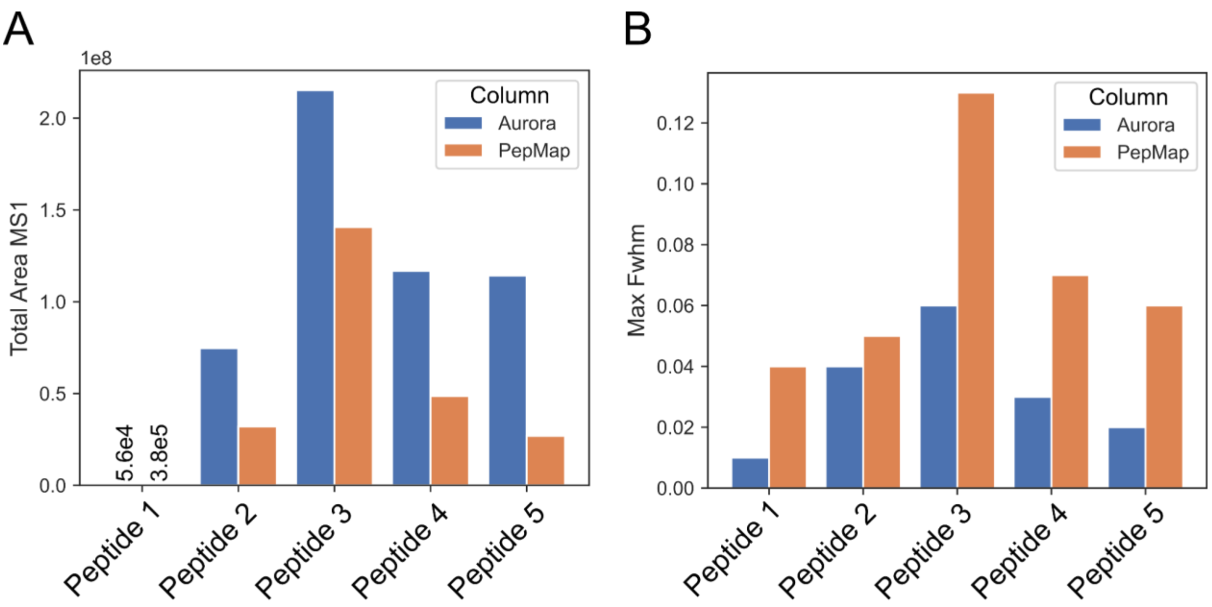

C

| | | |
|---|---|---|
| Peptide 1 | DKQSGK-GQKNSR | 19 min |
| Peptide 2 | LKGSPEDNEQK-ENQTTQKGQK | 23 min |
| Peptide 3 | KTEVQTGGFSKESILPK-KKDWDPK | 40 min |
| Peptide 4 | KSEETITPWNFEEVVDK-VNTEITKAPLSASMIK | 60 min |
| Peptide 5 | KSEETITPWNFEEVVDK-GILQTVKVVDELVK | 70 min |

*Figure S5: Column comparison of PepMap 50 cm vs. Aurora Ultimate 25 cm analytical columns using quantification of 5 crosslinked peptides across a 70 min gradient. A: Total MS1 Area of 5 selected crosslinked peptides, showing significantly higher peak areas for the Aurora column. B: Full-width half maximum times for each crosslinked peptide, showing broader peaks when using the PepMap column for crosslink samples. C: List of selected crosslinked peptides for quantitation in Skyline and their respective average retention times across the 70 min active gradient.*

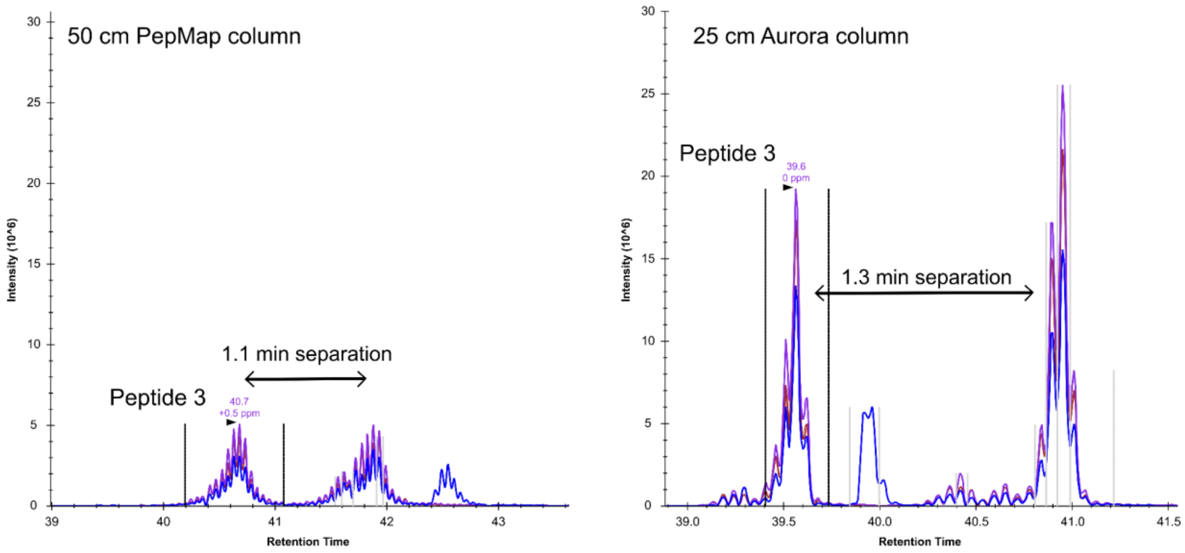

*Figure S6: Chromatographic separation comparison of PepMap 50 cm vs. Aurora Ultimate 25 cm analytical columns using chromatograms of peptide 3. Elution profiles were exported from Skyline (v 21.2.0.425) after peak*

*mapping and quantitation. PepMap chromatograms showing broadening and tailing of elution peaks of crosslinked peptide 3 and sharp well-separated peaks on the Aurora column.*

*Table S1: Column parameters for PepMap and Aurora analytical column used in this study.*

| Parameters | PepMap | Aurora |
|---|---|---|
| Length | 50 cm | 25 cm |
| Inner Diameter | 75 um | 75 um |
| Pore Size | 100 A | 120 A |
| Pressure Limit | 1500 bar | > 1700 bar |
| Max. Temperature Limit | 60°C | 60°C |
| Particle Size | 2 um | 1.7 um |
| Emitter | external fused silica emitter | Integrated |

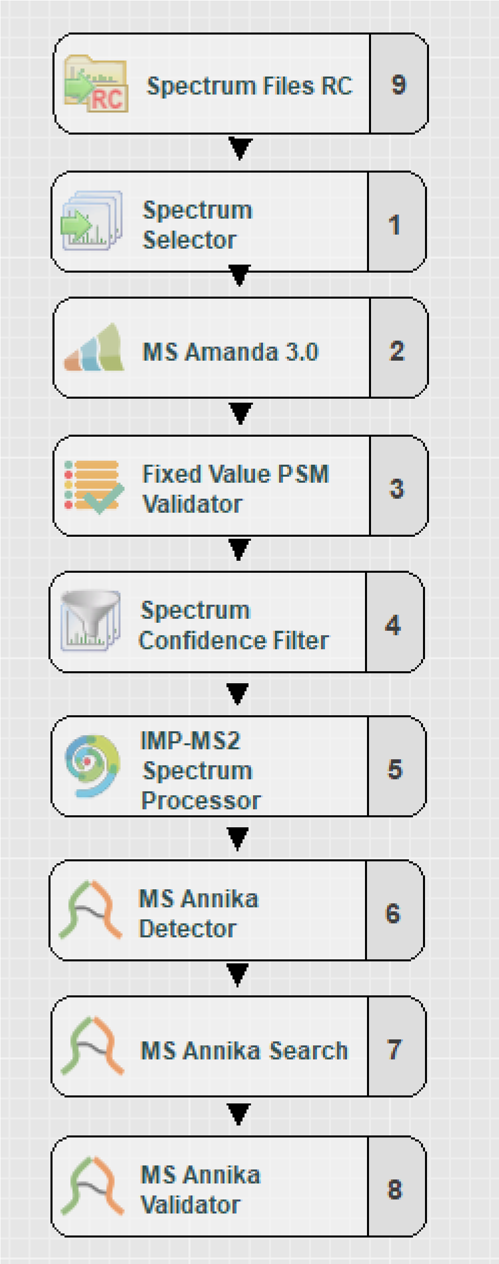

*Figure S7: Exemplary search workflow using MS Annika 3.0 in Proteome Discoverer. The imported raw files are recalibrated by the Spectrum file RC node, followed by a crude selection of suitable spectra for a first search by MS Amanda for linear peptide identification. Spectra that could not map to a linear or monolink peptide are transferred to the MS Annika crosslink search and validation nodes. The cross-link spectra are adjusted and then searched with MS Annika to identify the cross-linked peptides.*

*Table S2: Search parameters for linear and crosslink search. Parameters not listed here were left at default settings*

| *Parameter name* | *Parameter value* |
|---|---|
| ***Linear search*** | |
| *MS1 tolerance* | *6 ppm* |
| *MS2 tolerance* | *15 ppm* |
| *Miss cleavages* | *3* |
| *Fixed modification* | *Carbamidomethyl [57.021 Da] on C (D, E, H)* |
| *Variable modification* | *Oxidation [15.995 Da] (M),*<br><br>*PhoX [209.972 Da] (K), PhoX Amidated [226.998 Da] (K), PhoX Hydrolyzed [227.982 Da] (K), PhoX Tris [331.046 Da] (K)*<br><br>*DSSO [158.004 Da] (K), DSSO Amidated [175.030 Da] (K), DSSO Hydrolyzed [176.014 Da] (K), DSSO Tris [279.078 Da] (K)* |
| ***Crosslink search*** | |
| *PhoX*<br>*DSSO* | *PhoX [209.972 Da] (K)*<br>*DSSO [158.004 Da] (K) (additional doublet: 49.982635 Da)* |
| *MS1 tolerance* | *6 ppm* |
| *MS2 tolerance* | *15 ppm* |
| *Miss cleavages* | *3* |
| *Fixed modification* | *Carbamidomethyl [57.021 Da] (C)* |
| *Variable modification* | *Oxidation [15.995 Da] (M)* |
| *Top N* | *500* |

| | |
|---|---|
| *Candidate Search Method* | *i32SM* |
| ***IMP-MS2 spectrum processor*** | |
| *Perform de-isotoping* | *False* |

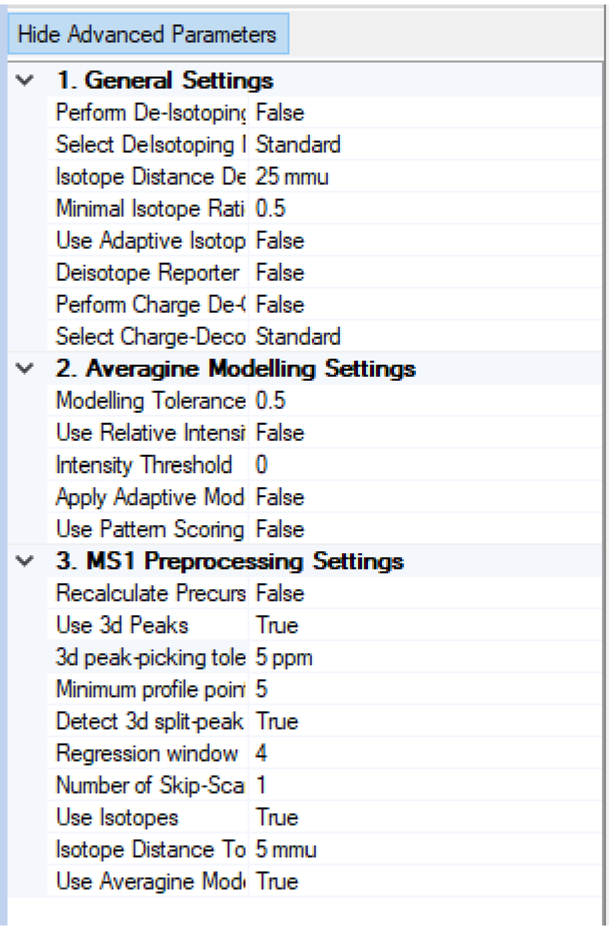

*Figure S8: General settings of the IMP-MS2 Spectrum processor node. This node offers three spectrum processing steps: Deisotoping of isotopic clusters, Charge-Deconvolution and MS1-precursor recalculation. The algorithm reconstructs the elution profile of the peptide and uses the gathered data to calculate a more precise value of the precursor mass. This step was evaluated as beneficial for crosslinking data.*

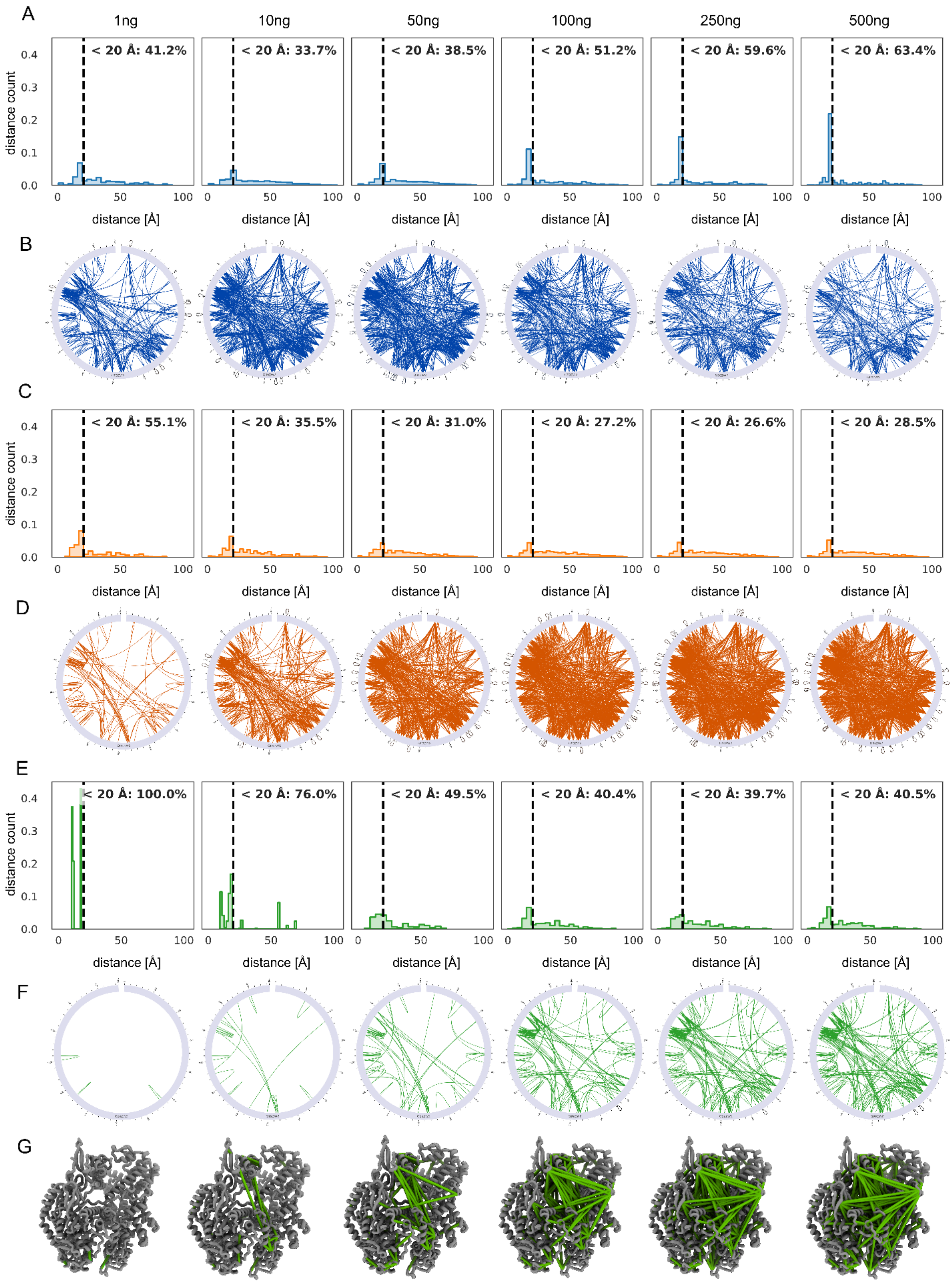

*Figure S9: Evaluation of the biological relevance of crosslinks across low, medium, and high MS1 precursor intensities. Crosslinks were categorized based on MS1 apex intensity into low ($< 1 \times 10^5$), medium ($\geq 1 \times 10^5$ and $< 1 \times 10^7$), and high ($> 1 \times 10^7$) abundance groups, and analyzed across a dilution series from 1 ng to 500 ng of Cas9 crosslinked with PhoX. A: Distribution of crosslink distances for low-abundance precursors (blue), mapped onto an AlphaFold3-predicted Cas9 structure. The percentage of crosslinks falling below the 20 Å structural distance threshold increases with higher injection amounts, suggesting greater structural compatibility at*

*improved detection levels. B: xiView circular representation of low-abundance (< 1 × 10⁵) crosslinks for Cas9. The total number of detected links increases with injection amount, but no clear pattern of biologically preferred linkage regions is observed. C: Distribution of crosslink distances for medium-abundance precursors (orange). In contrast to panel A, the percentage of links below 20 Å decreases as injection amounts increase, suggesting a growing presence of structurally non-ideal or potentially inter-molecular links. D: xiView circular view of medium-intensity crosslinks (1 × 10⁵ to 1 × 10⁷). Link numbers increase with sample amount but remain broadly distributed across the protein. E: Distribution of crosslink distances for high-abundance precursors (green). The proportion of short-range crosslinks (< 20 Å) decreases with increasing injection, though all high-abundance links at 1 ng fall within the 20 Å limit, indicating structurally consistent and likely intra-molecular links at low sample loads. F: xiView circular view of high-intensity (> 1 × 10⁷) crosslinks. As with other groups, link numbers increase with higher injection amounts, but no specific structural bias is evident. G: 3D structural representation of Cas9 with high-intensity crosslinks mapped. At 1 ng injection, crosslinks are restricted to surface-accessible regions, while at higher injection amounts, internal regions of the protein become increasingly covered, suggesting deeper penetration into protein structure with increasing precursor abundance.*

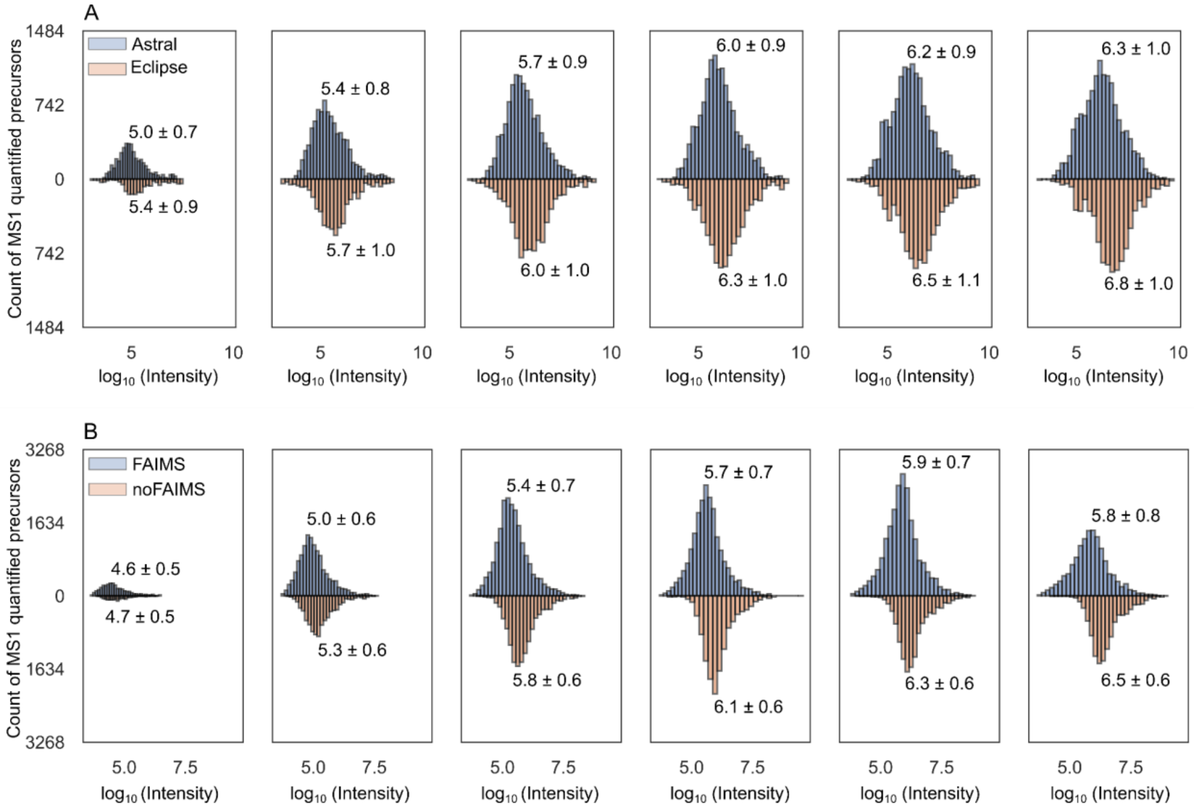

*Figure S10: Abundance distribution of MS1-quantified precursors comparing Orbitrap Astral vs. Eclipse and FAIMS vs. no FAIMS measurements. A: Distribution of MS1-quantified precursor intensities across a dilution series from 1 ng to 500 ng, comparing Orbitrap Astral (blue) and Orbitrap Eclipse (orange). For each condition, medians and standard deviations (calculated as the average distance of individual values to the sample mean) are indicated above the histograms. Across all injection amounts, Astral measurements consistently exhibit lower median intensities, reflecting the instrument's increased sensitivity relative to Eclipse. B: Distribution of MS1-quantified precursor intensities comparing Orbitrap Astral with FAIMS (blue) and without FAIMS (orange) across the same dilution series. Application of FAIMS results in lower median precursor intensities across all injections, attributed to reduced chemical noise and enhanced sensitivity through gas-phase ion filtering.*

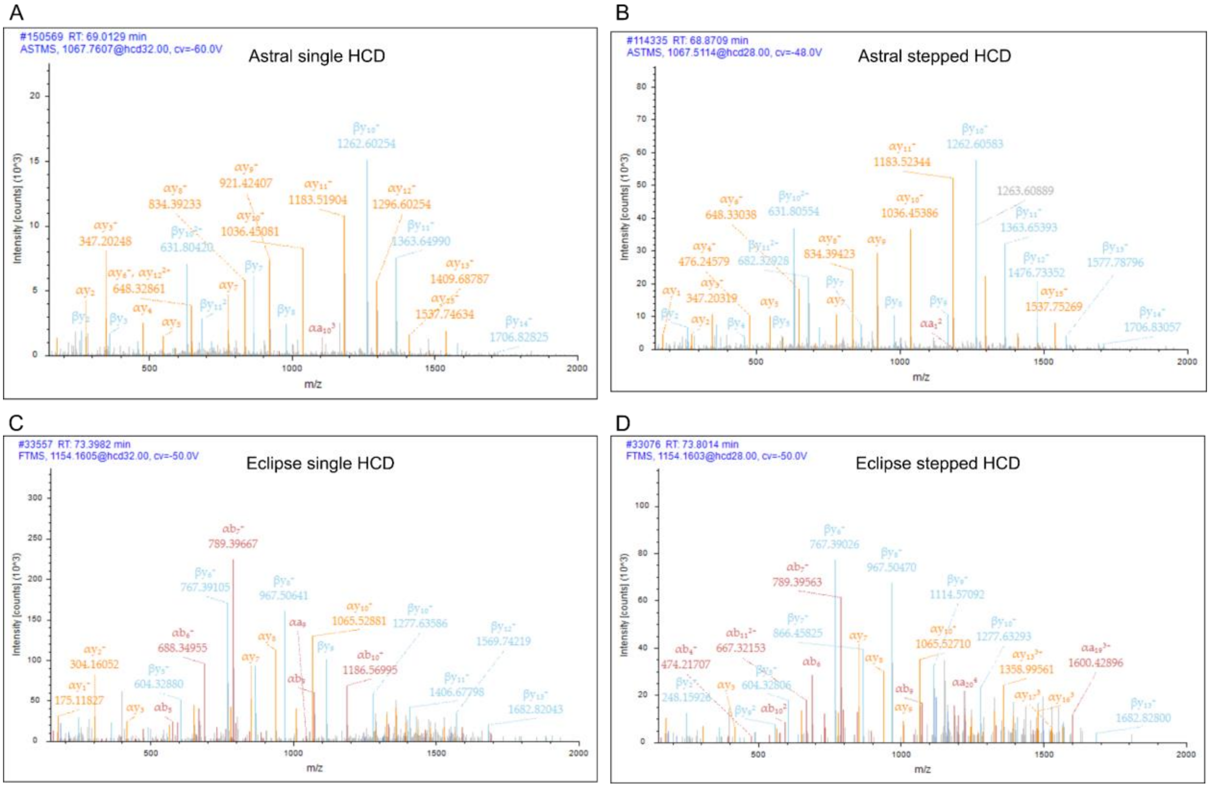

*Figure S11: Representative MS2 fragmentation spectra for crosslinked peptides using single and stepped HCD on Astral and Eclipse instruments. A: Peptide KNLIGALLFDSGETAEATR–KSEETITPWNFEEVVDK with charge 4 on RT 69 min acquired on Orbitrap Astral using single HCD. B: Same peptide acquired on Astral using stepped HCD. C: Peptide KSEETITPWNFEEVVDKGASAQSFIER–VLPKHSLLYEYFTVYNELTK with charge 5 on RT 73 min acquired on Orbitrap Eclipse using single HCD. D: Same peptide using stepped HCD. Fragmentation spectra of stepped HCD shows slightly improved fragmentation of higher m/z fragments, though both fragmentation methods yield high-quality MS2 spectra overall for both instruments.*

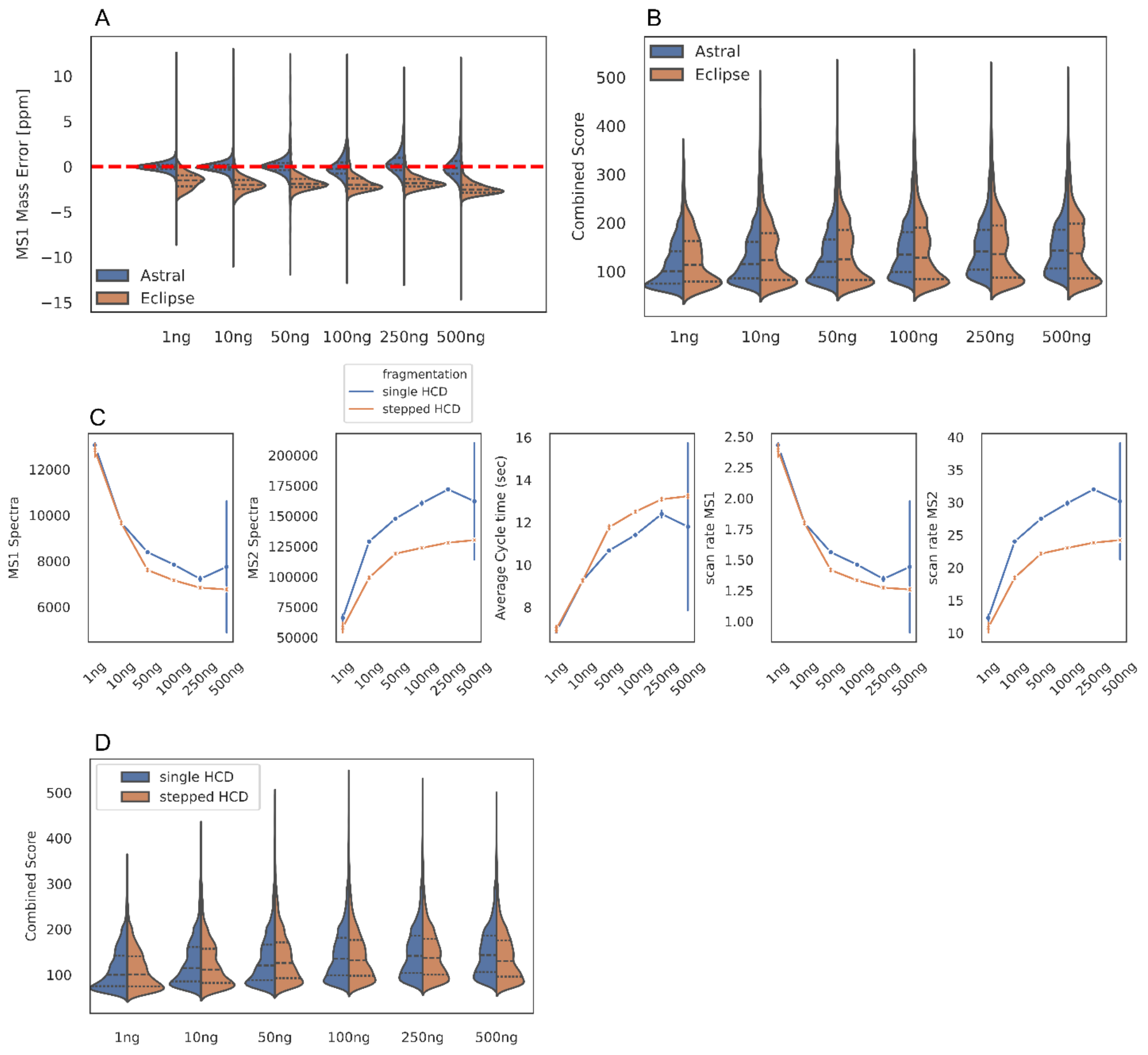

*Figure S12: MS1 mass error and score distributions for PhoX-crosslinked Cas9 across injection amounts (1–500 ng). A: MS1 mass error distributions for Orbitrap Astral and Eclipse acquired using identical parameters (500 AGC target, 6 ms injection time). Astral data exhibit well-centered mass errors across all injection amounts, while Eclipse data show slightly off-centered distributions. B: Distribution of combined scores for the PhoX dilution series (1–500 ng). Astral data yield slightly higher scores than Eclipse at higher injection amounts (100–500 ng), whereas scores are marginally lower for Astral at low injection amounts (1–50 ng). C: MS1 scans, MS2 scans, average cycle time, MS1 scan rate (IT: 6ms) and MS2 scan rate (IT: 20ms) for single HCD and stepped HCD methods on the Orbitrap Astral. MS1 and MS2 scans are only increased for single HCD, while average cycle time is faster for single HCD methods. Scan rates for MS1 and MS2 also benefit from single HCD methods with faster scan rates for MS2 specifically. D: Distribution of combined scores for the PhoX dilution series (1–500 ng) for single and stepped HCD, showing slightly better score medians across conditions for single HCD. Note, that MS Annika does perform preprocessing of the spectra before searching crosslinked peptide. This likely removes any "noise" introduced by the higher sensitivity of the Astral analyzer.*
